# Crossing-over decision landscape in maize

**DOI:** 10.1101/2022.09.21.508771

**Authors:** Mateusz Zelkowski, Minghui Wang, Qi Sun, Jaroslaw Pillardy, Penny M.A. Kianian, Shahryar F. Kianian, Changbin Chen, Wojciech P. Pawlowski

**Affiliations:** School of Integrative Plant Science, Cornell University, Ithaca, NY 14853, USA; Bioinformatics Facility, Cornell University, Ithaca, NY 14853, USA; Department of Horticultural Science, University of Minnesota, St. Paul, MN 55108, USA; USDA-ARS, Cereal Disease Laboratory, St. Paul, MN 55108, USA; School of Life Sciences, Arizona State University, Tempe, AZ 85287, USA

## Abstract

In most crops, including maize, meiotic double-strand breaks (DSBs) occur in all chromosome regions but crossovers (COs) are predominantly near chromosome ends. To understand how the uniform DSB distribution changes into the U-shaped CO distribution, we generated high-resolution maps of CO intermediates. We found that DSBs with medium resection spans more often result in COs than those with shorter or longer resections. We also discovered that sites of CO intermediates associated with MLH3 in zygotene are uniformly distributed along chromosomes, resembling DSB distribution. However, in late prophase, they show the U-shaped distribution characteristic of COs. While zygotene MLH3 sites exhibit methylation levels similar to the genome average, late prophase sites have reduced DNA methylation. In contrast to DNA methylation, inter-parental DNA sequence polymorphism has limited effect on CO distribution. These data indicate that the final CO landscape shape in maize is established late during recombination and controlled by chromatin state.

## INTRODUCTION

The main source of genetic diversity in most eukaryotes are DNA strand exchanges between homologous chromosomes called crossing-overs (COs) that are generated during meiosis. Despite their importance, COs are not evenly spaced along chromosomes and tend to form distinct hotspots (Zelkowski et al., 2019). Furthermore, plants with large genomes, such as maize or wheat, exhibit a strong bias in CO distribution, with the majority of them found near chromosome ends and extensive regions surrounding the centromeres exhibiting very few COs (Gore et al., 2009; Kianian et al., 2018; Zelkowski et al., 2019). This distribution pattern has profound consequences for evolution (Yang et al., 2017) and breeding (Taagen et al., 2020).

COs are products of meiotic recombination, which starts with the programed formation of double-strand breaks (DSBs) in chromosomal DNA induced by the SPO11 protein complex (Bergerat et al., 1997; Ku et al., 2020). Resection of ends of these DSB ends leads to the formation of single-stranded DNA (ssDNA) overhangs (Garcia et al., 2011; Lukaszewicz et al., 2015). The overhangs become substrates for RAD51- and DMC1-mediated single-end invasion of homologous chromosomes (Savocco and Piazza, 2021). This process eventually creates double Holliday junctions (dHJs), which can lead to the formation of COs (Schwacha and Kleckner, 1995). However, relatively few DSBs result in COs. In maize, there are around 500 DSBs per meiocyte, yielding approximately 18 COs (Sidhu et al., 2015). In contrast to the biased CO distribution (Kianian et al., 2018), DSBs in maize are found chromosome-wide, including centromeric and pericentromeric regions (He et al., 2017).

Most COs, at least 85% of them in maize (Falque et al., 2009), are products of the class I CO pathway, which is facilitated by the group of ZMM proteins (Pyatnitskaya et al., 2019). Key members of this group are MutLγ proteins MLH1 and MLH3 (Franklin et al., 2006; Hunter and Borts, 1997; Xin et al., 2021). These two proteins form a heterodimer, which processes dHJ intermediates (Manhart et al., 2017). Down-regulation of MutLγ proteins results in drastic CO number reduction in a variety of organisms (Colas et al., 2016; Jackson et al., 2006; Mao et al., 2021; Nishant et al., 2008; Svetlanov et al., 2008).

The decision between CO and non-CO products remains enigmatic at the molecular level. DSBs may be marked to become COs very early in meiosis (Borner et al., 2004). In maize, the sites of the DSBs that are more likely to result in COs exhibit distinct chromatin features from the DSB sites where COs are rarely formed, primarily with regard to the DNA methylation level (He et al., 2017). However, the final CO/non-CO decision may be executed late in meiosis. Cytological studies in wheat and barley have shown that numbers of MLH3 foci on chromosomes decrease during prophase I (Colas et al., 2016; Osman et al., 2021), suggesting that MLH1/3 are associated not exclusively with the dHJs resulting in COs.

To elucidate the dynamics of CO formation, we investigated two key steps in the CO pathway. From studying DSB resection, we found that DSBs with intermediate resection lengths are more likely to result in COs. To examine late steps of CO formation, we created high-resolution maps of chromosomal sites of MLH3 throughout prophase I using chromatin immuno-precipitation (ChIP). We discovered that MLH3 distribution in early prophase I mimics DSB distribution while MLH3 distribution in late prophase resembles CO distribution. This observation implicates MLH3 in the final step of the CO/non-CO decision. We furthermore found that this decision step is tightly associated with DNA methylation levels at dHJ sites. Finally, we also demonstrate that MLH3 ChIP is a fast, high-throughput, and efficient method to generate genetic maps in diverse germplasm, including backgrounds with low levels of inter-parental DNA sequence polymorphism.

## RESULTS

### DSB sites with 500 - 1500bp resection spans are more likely to result in COs

To elucidate the exact steps of meiotic recombination at which the CO/non-CO decision takes place, we first conducted an analysis of meiotic DSB resection. We hypothesized that the span of ssDNA resection may affect the likelihood of a break being repaired as a CO vs. a non-CO. To examine DSB resection, we performed S1 nuclease mapping (Mimitou et al., 2017) on zygotene-stage anthers in the maize inbred B73. Sequencing of Illumina libraries of DNA fragments produced by S1 treatment resulted in 6.8mln and 11.9mln reads (Supplemental Table 1) in two highly similar biological replicates (Spearman rank correlation coefficient = 0.93) (Supplemental Table 2). In contrast, the negative control generated under the same conditions resulted in only 1.8mln reads for an untreated zygotene sample, 6766 reads for an S1-treated leaf sample, and 5168 reads for an S1-treated sample of anthers from the *ameiotic1-489* (*am1-489*) mutant (Supplemental Table 1), which lacks meiotic DSBs (Pawlowski et al., 2009).

To determine the span of DSB resection, we measured the distance between adjacent peaks formed when Illumina reads are aligned to the genome, which should represent opposing ends of DSBs. Compared to the untreated control and random reads from genomic DNA, we observed in the S1-treated zygotene samples a significant (*P* < 2.2e^-16^, Chi-square test) enrichment of peaks located 500 to 2000bp apart, with 1079bp being the median distance (Figure 1A). This value is similar to the ∼0.8kb and 1.1kb DSB resection spans reported in yeast and mouse, respectively (Marsolier-Kergoat et al., 2018; Yamada et al., 2020). Of the 9799 maize DSB hotspots identified using RAD51 ChIP (He et al., 2017), 53.3% were located within 5kb from one or more of the roughly 50,000 S1 resection sites 500bp to 2000bp in length (*P* < 2.20e^-16^ compared to random genome sites) (Supplemental Figure 1).

**Figure 1.**
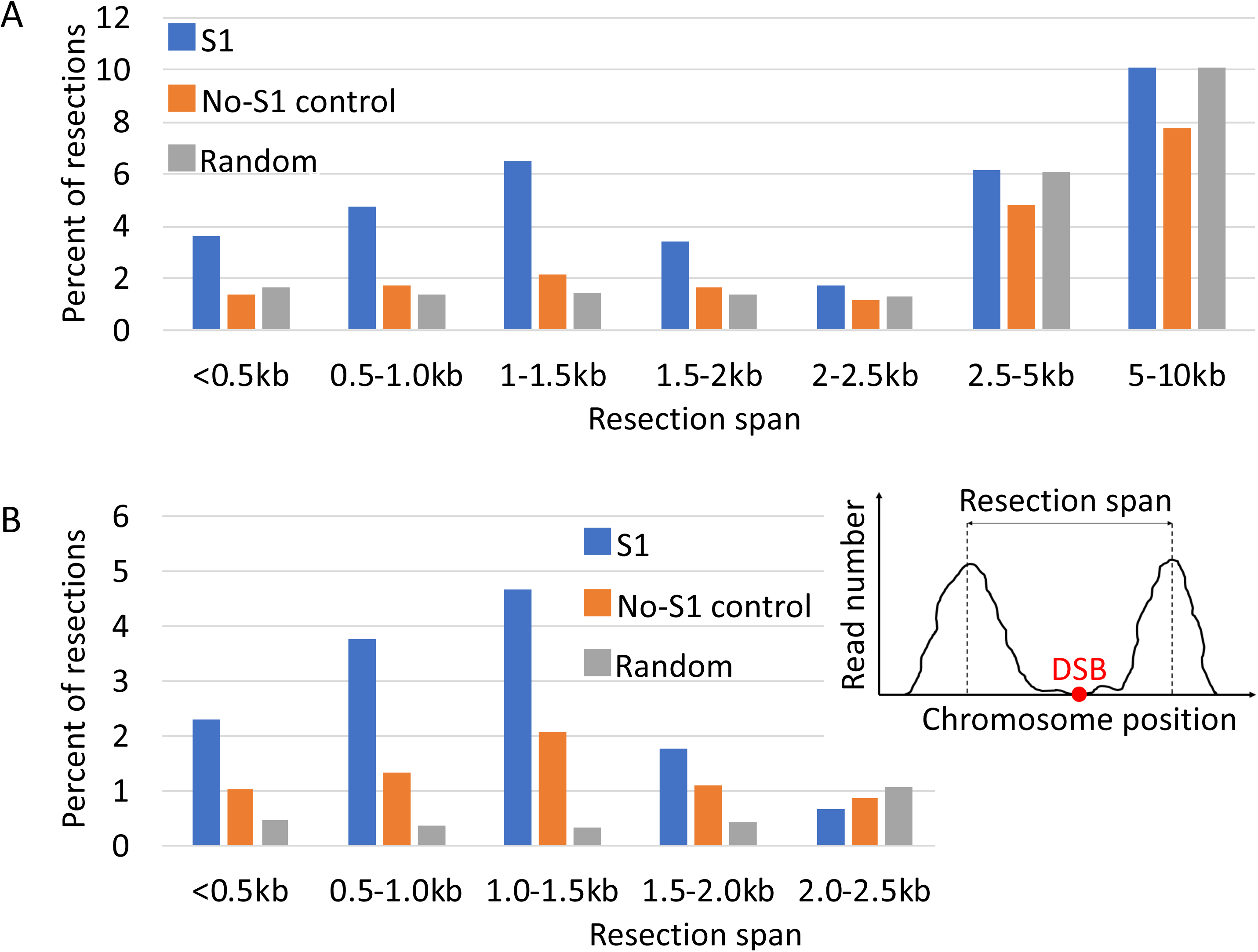
Analysis of DSB resection in maize using S1 nuclease mapping. S1 = B73 zygotene anthers treated with S1 nuclease, Random genome sites, No S1 treatment = B73 zygotene anthers without S1 treatment. **B.** Frequency of NAM COs located within 5kb from DSB resection sites of different lengths.

To determine whether a specific resection span was more likely to result in CO formation, we related locations of S1 peak pairs to the positions of ca. 75,000 COs identified in the maize Nested Association Mapping (NAM) population produced by crossing B73 to 25 diverse inbreds (McMullen et al., 2009). Roughly 60% of these COs were located within 10kb from resection sites 500bp to 1500bp in span (*P* < 2.2e^-16^ compared to random genome sites). Interestingly, resections sized 500bp to 1500bp, were more likely to colocalize with COs than resections with shorter or longer spans (Figure 1B).

### MLH3 chromosomal foci decrease in number during prophase I progression

To study the dynamics of CO formation at the dHJ resolution stage, we produced two antibodies, targeting the N- and C-termini of MLH3. Microscopic evaluation showed that the antibodies co-localized with a high degree, identifying distinct foci on meiotic chromosomes (Supplemental Figure 2). The foci first appear in zygotene and persist until diplotene (Figure 2A). Their numbers decrease during prophase progression, starting from 30.2±8.1 (n=29) in zygotene to 25.3±5.4 (n=10) in pachytene and 13.0±1.7 (n=6) in diplotene. Some but relatively few of the zygotene MLH3 foci colocalized with DSB markers RAD51 and γH2AX (Supplemental Figure 3). The MLH3 focus count in diplotene corresponded roughly to the number of class I COs in maize (Sidhu et al., 2015 2009). These data suggest that zygotene and pachytene MLH3 sites correspond to CO intermediates whereas diplotene MLH3 sites represent class I COs.

**Figure 2.**
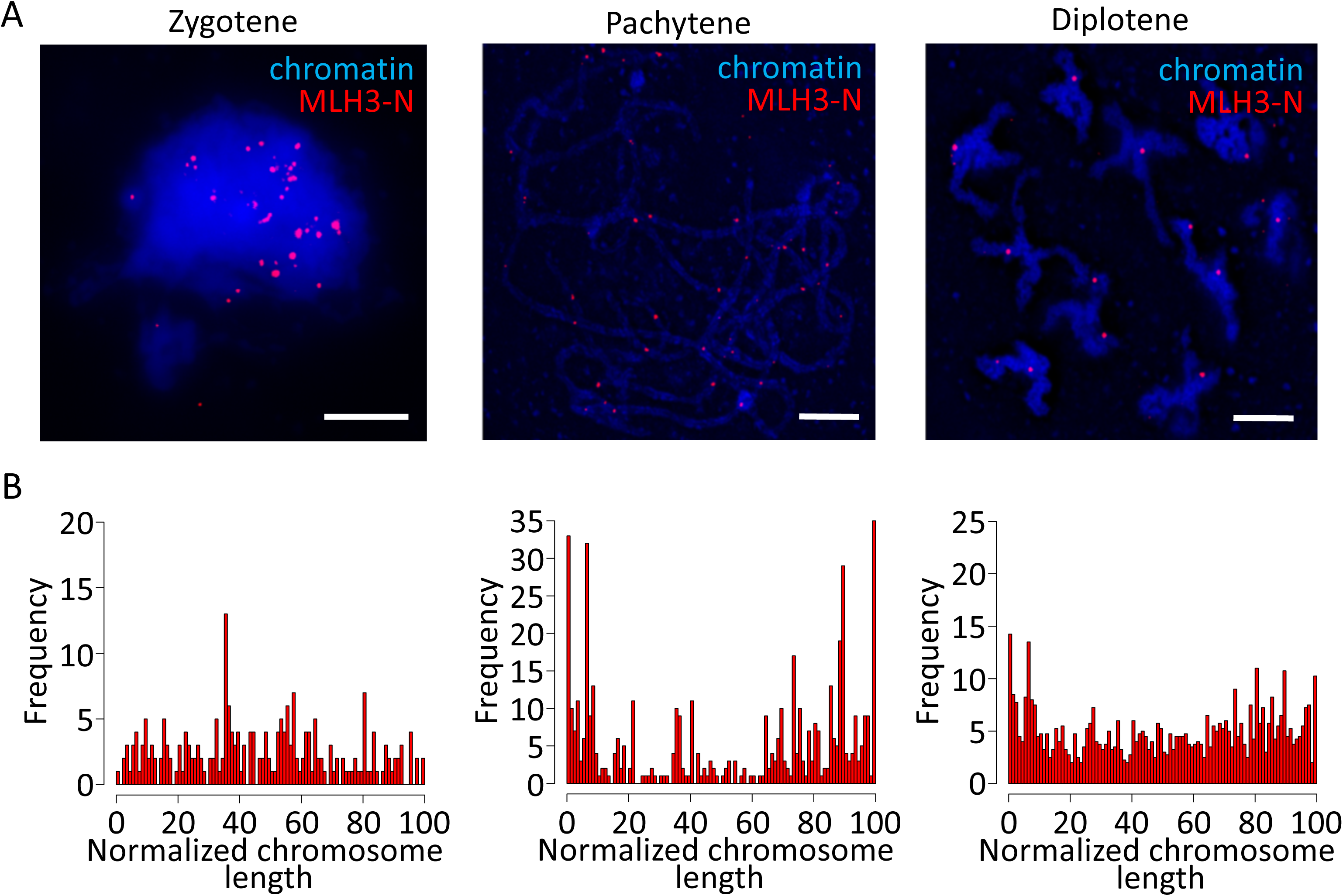
Distribution of MLH3 during prophase I. **A.** Immunolabelling of MLH3 foci using the MLH3-N antibody in meiocytes of the B73 x Mo17 hybrid. Scale bar = 5μm. **B.** Distribution MLH3-N peaks along B73xMo17 maize chromosomes. Figures are summaries of all ten chromosomes normalized to the same length.

### ChIP-identified MLH3 hotspots exhibit distinct distribution in early and late prophase I

After the cytological validation, we used the anti-MLH3 antibodies in chromatin immunoprecipitation (ChIP) experiments on zygotene, pachytene, and diplotene maize tassels from the maize B73 x Mo17 hybrid. In these experiments, chromatin fragments associated with MLH3 were purified, and the corresponding DNAs were sequenced and mapped to the maize genome (Supplemental Table 3). Consistently with the immuno-localization results, results of ChIP-seq experiments using the antibodies directed at the N- and C-termini of MLH3 were highly correlated to each other (Supplemental Table 4).

Analysis of the sequence data revealed presence of distinct hotspots in all three prophase I substages. Increasingly more hotspots were present with the progression of prophase: the average of MLH3-N and MLH3-C experiments being 375 hotspots in zygotene, 1090 hotspots in pachytene and 35,185 hotspots in diplotene. Positions of pachytene and diplotene hotspots were highly similar to each other, with up to 90.3% of pachytene hotspots colocalizing with diplotene hotspots. In contrast, zygotene hotpots were quite different, with only 2% to 27.9% of them colocalizing with late prophase (pachytene and diplotene) hotspots (Supplemental Table 5). Overall, these results revealed two classes of MLH3 hotspots, early-prophase hotspots and mid/late-prophase hotspots.

Numbers of MLH3 hotspots per chromosome were generally proportional to chromosome length. However, hotspot distribution patterns were distinct between early and late prophase substages. In zygotene, MLH3 hotspots were distributed fairly uniformly along chromosomes, exhibiting similar hotspot density and strength in sub-telomeric as in pericentromeric chromosome regions (Figure 2B). These distribution patterns resembled the distribution of meiotic DSBs (He et al., 2017). In contrast, MLH3 hotspots in pachytene and diplotene showed U-shaped distribution, with elevated numbers in distal chromosome region (Figure 2B), which is characteristic of CO distribution in maize (Kianian et al., 2018).

### Late prophase MLH3 hotspots colocalize with genetically-mapped COs

To further investigate MLH3 hotspot distribution, we compared hotspots from the three prophase substages to the 75,000 NAM population COs (McMullen et al., 2009). MLH3 hotspots identified in zygotene did not show an overall overlap with known COs (window size = 10kb) at a level significantly higher than random genome sites (*P* = 1). In contrast, we found statistically significant colocalization between the sites of genetically-mapped CO and late prophase MLH3 hotspots (*P* = 3.4e^-8^ for pachytene MLH3 hotspots and *P* = 0.073 for diplotene hotspots). Among late-prophase MLH3 hotspots, 25% were in gene bodies, 15% were within 2kb downstream from transcription termination sites, 10% were within 2kb upstream from transcription start sites, and 50% were further than 2kb away from genes (Figure 3). These values were similar to those characterizing genetically-mapped COs (Kianian et al., 2018).

**Figure 3.**
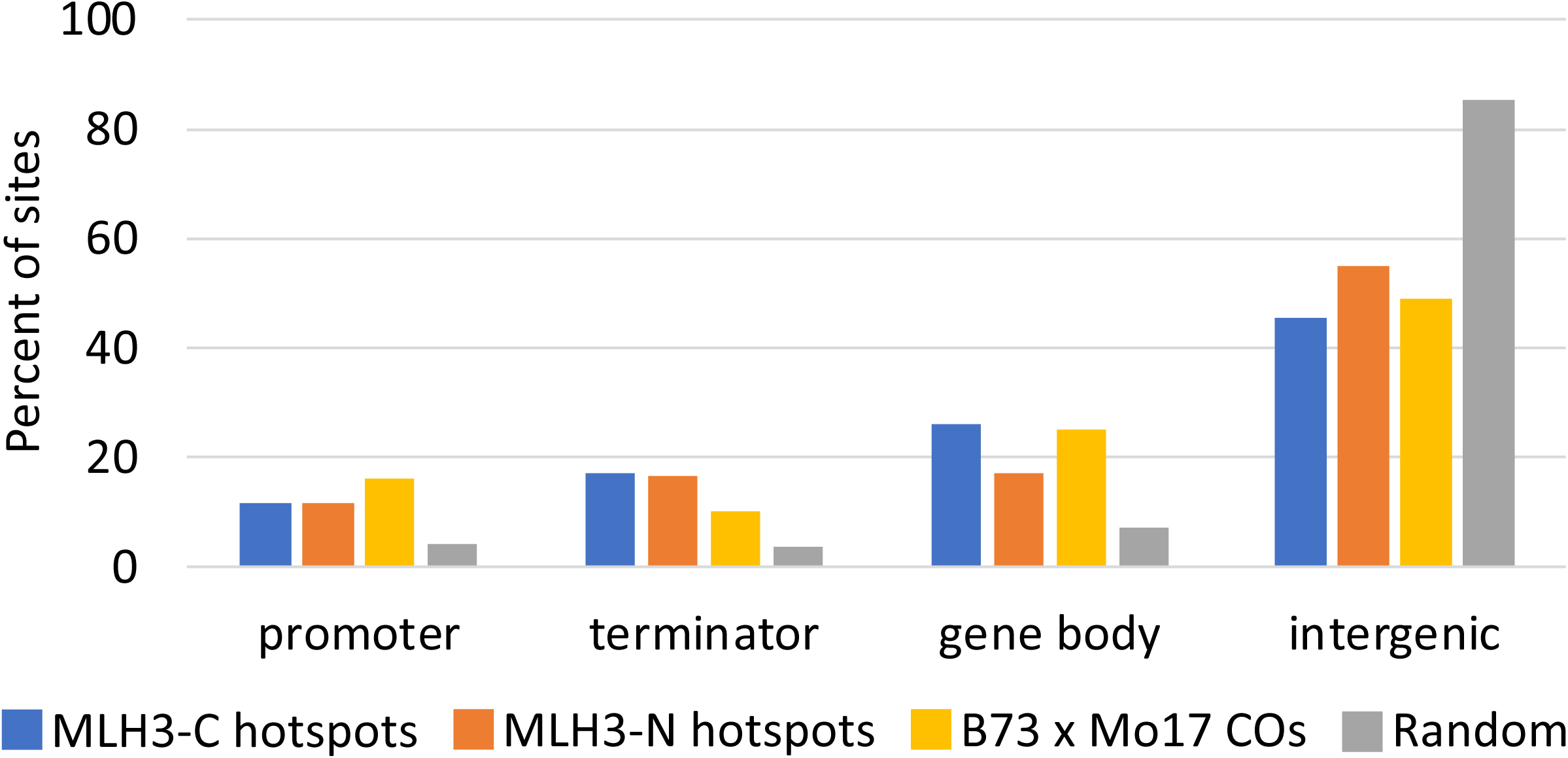
Localization of diplotene MLH3 hotspots in different gene parts in the B73 x Mo17 hybrid compared to NAM population COs.

### A large portion of MLH3 hotspots are in single-nucleotide polymorphism deserts

Substantial segments of the maize genome exhibit limited sequence polymorphism, most likely stemming from selective sweep episodes during maize evolution and domestication (Tian et al., 2009). These polymorphism deserts reduce the resolution of genetic mapping of COs. In contrast to genetic mapping, MLH3 ChIP provides high resolution regardless of polymorphism. About 22% of the B73 inbred genome are segments 100kbp or more that lack single nucleotide polymorphisms (SNP) compared to the Mo17 inbred genome. We found that roughly 40% of maize of pachytene and diplotene MLH3 hotspots were in these genomic regions.

### MLH3 hotspot landscape in a homozygous line is similar to that of a hybrid

Several studies in maize and other plants have suggested that DNA sequence polymorphism exhibits a major effect on CO landscape (Blackwell et al., 2020; Dooner, 2002; Emmanuel et al., 2006; Ziolkowski et al., 2015). As MLH3 ChIP can be used to create CO maps in homozygous backgrounds, this question can now be addressed on the genome-wide scale. To do this, we generated an MLH3 hotspot map in the maize inbred B73. Overall, MLH3 hotspot patterns in B73 mimicked those of the B73 x Mo17 hybrid at all three prophase substages examined (Figure 4A). We also found strong correlation between the positions of diplotene MLH3 hotspots in the B73 inbred and the B73 x Mo17 hybrid at finer scales (*r* = 0.88 using 10kb window size). Furthermore, 39.7% of MLH3 hotspots in diplotene were located within 5kb from COs genetically-mapped in the maize NAM population.

**Figure 4.**
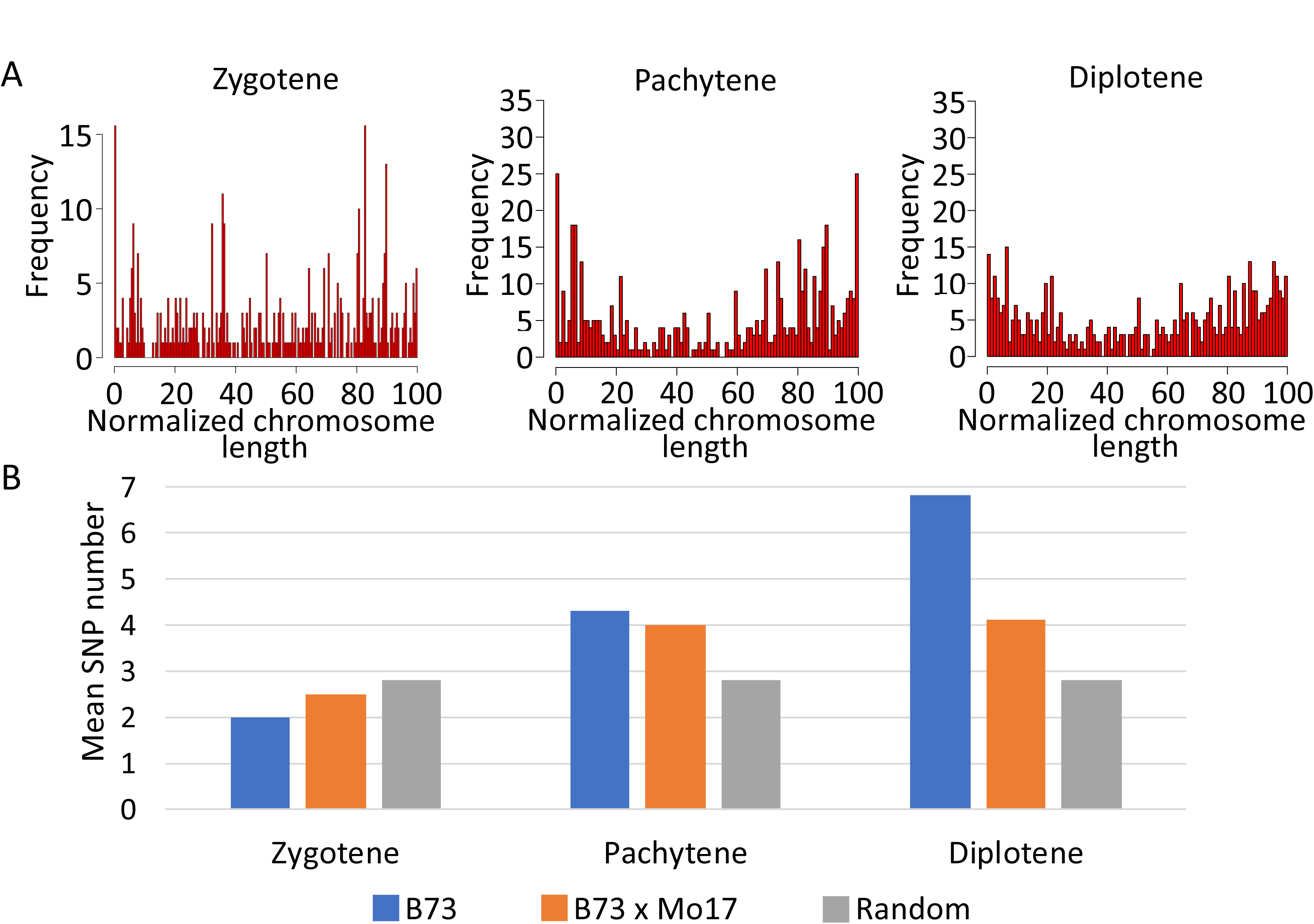
Distribution of MLH3 peaks in the B73 maize inbred line. **A.** Distribution MLH3-N peaks along B73xMo17 maize chromosomes. Figures are summaries of all ten chromosomes normalized to the same length. **B.** Mean number of SNPs between the B73 x Mo17 genomes in 2kb-wide regions centered on MLH3 hotspots identified in the B73 inbred and the B73 x Mo17 hybrid.

In addition, we examined SNP levels at MLH3 hotspots identified in the B73 x Mo17 hybrid. We found that the sites of pachytene and diplotene hotspots exhibited higher SNP levels than the sites of zygotene hotspots (Figure 4B). Comparing MLH3 hotspot sites of the hybrid to those of the B73 inbred, we discovered that the zygotene and pachytene hotspots in B73 corresponded to locations exhibiting SNP levels similar to the zygotene and pachytene hotspots of the hybrid (Figure 4B). In contrast, diplotene hotspots of the hybrid were at sites exhibiting much lower SNP levels than the sites corresponding to the diplotene hotspot sites of B73. These data imply preferential CO formation at sites with lower SNP levels.

### MLH3 hotspots display conserved DNA sequence motifs and are frequently located at chromosome knobs

The high resolution of ChIP-seq affords a unique ability to examine the precise DNA sequence context of CO and CO intermediate sites. To exploit this opportunity, we conducted a detailed analysis of 500 strongest MLH3 hotspots from each of the three prophase I substages. We found that in the B73 x Mo17 hybrid, 68.9%, 50.0% and 28.4% of zygotene, pachytene and diplotene hotspot sites, respectively, displayed conserved DNA sequence motifs (*P* < 1e^-10^) (Figure 5A-D). These motifs were located at an average of 267bp from MLH3 hotspot centers. The most common motif (Figure 5A), present at 28% of the 500 strongest hotspots in diplotene and at 10% of hotspots in pachytene, was part of the 180bp repeat found in maize heterochromatic knobs (Peacock et al., 1981). However, not all of the 180bp repeat-containing knobs present in the B73 and Mo17 genomes were MLH3 hotspots (Figure 5E). Similar observations, showing that only some knobs are CO sites, were made in a study of genetically-mapped COs (Kianian et al., 2018) and in immunocytology analyses (Stack et al., 2017). We confirmed localization of MLH3 foci at chromosome knobs with immunolocalization microscopy (Figure 5F).

**Figure 5.**
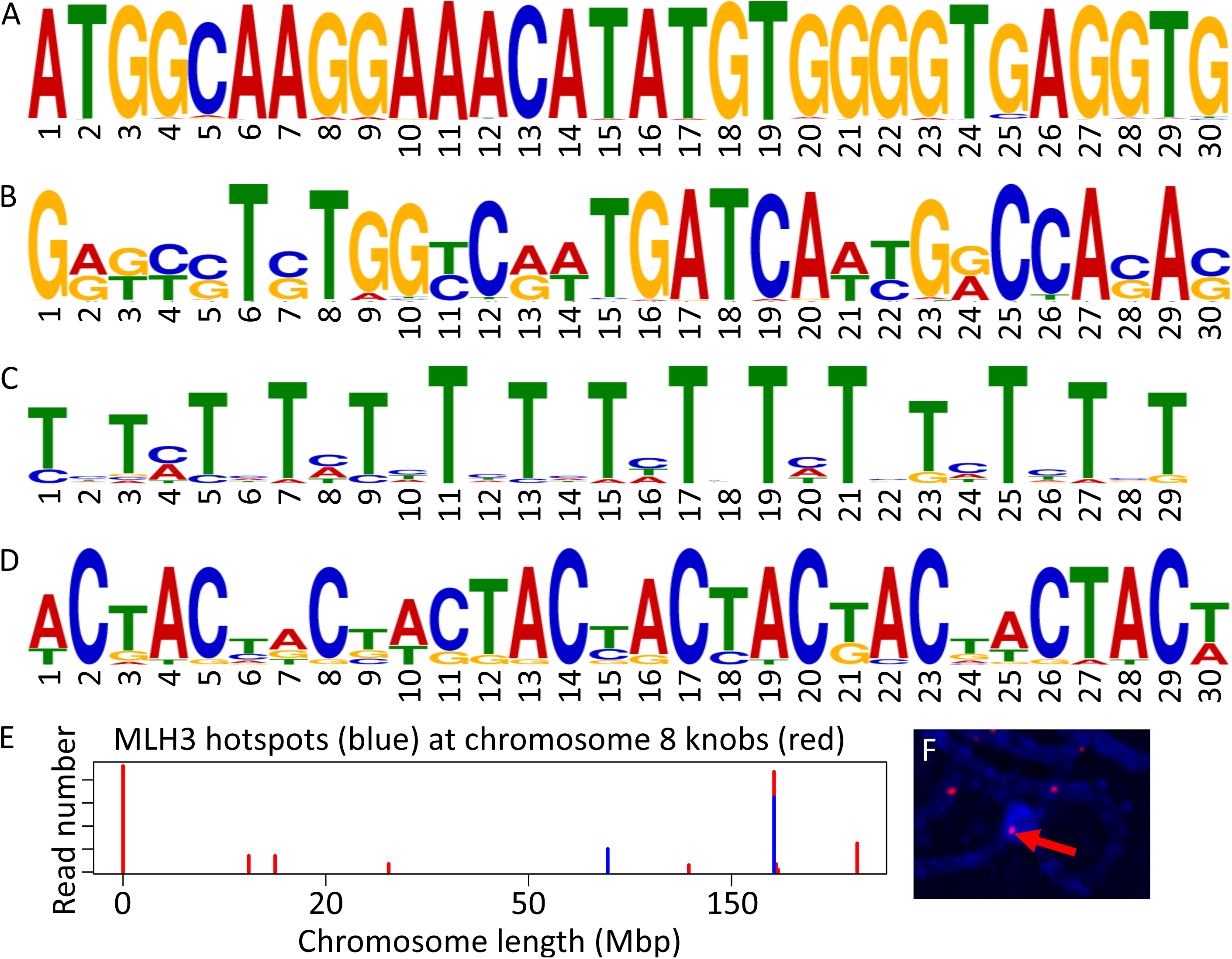
DNA sequence motifs at MLH3 hotspots. **A - D.** Four most common DNA sequence motifs. **A.** Highly conserved MLH3 hotspot motif that is part the 180bp knob repeat. **B.** Conserved motif that is also part of the 180bp knob repeat and displays a palindromic sequence at nucleotide positions 6 to 29. **C.** Thymidine-enriched motif. **D.** Cytidine-enriched motif. **E.** Position and number of the 180bp knob repeats on chromosome 8 relative to the position and strength of knob motif-localizing MLH3 hotspots. **F.** Antibody staining of MLH3 sites on late diplotene chromosomes in B73 showing an MLH3 focus at a DAPI-stained knob structure.

Other motifs associated with the 500 strongest MLH3 hotspots included an A-rich polymer and a degenerate C-rich trinucleotide repeat, as well as a novel 20-nt long palindrome (Figure 5C and D). A-rich polymers have been previously identified at CO sites in maize and Arabidopsis (Choi et al., 2013; Kianian et al., 2018; Shilo et al., 2015; Wijnker et al., 2013) and are thought to be associated with low nucleosome occupancy (Kaplan et al., 2009). Palindromic motifs have been reported at SPO11 and TOP2 cleavage sites in yeast (Gittens et al., 2019; Prieler et al., 2021).

### Late prophase MLH3 hotspots are closer to CO sites than early prophase hotspots

To study the dynamics of resolution of CO intermediates into COs, we calculated the distance between MLH3 hotspots and CO sites. For this analysis, we used ∼11,000 NAM population COs that were mapped with the resolution of 10kb or less, and only considered MLH3 hotspots whose center was no further than 10kb from CO midpoint. We found that the average distance from CO sites was 8550bp for zygotene MLH3 hotspots, 4292bp for pachytene hotspots and 3363bp for diplotene hotspots in B73 and 9962bp, 5863bp, and 6601bp, respectively, in the B73 x Mo17 hybrid. These observations show that MLH3 sites become closer to the eventual CO sites with the progression of recombination.

### Early and late MLH3 hotspots exhibit distinct DNA methylation patterns

To dissect the differences in MLH3 hotspot localization patterns in early *versus* late prophase, we examined chromatin characteristics of hotspot sites. We found that cytosine methylation in the CG and CHG contexts (where H is any nucleotide other than C) was reduced at MLH3 hotspots compared to the genome average (Figure 6). In contrast, CHH methylation was elevated. Interestingly, the differences between DNA methylation levels of MLH3 hotspots and the genome average were increasingly more pronounced at late prophase hotspots, showing a clear progression from zygotene to pachytene and to diplotene (Figure 6). Nucleosome occupancy levels, measured by monococcal nuclease sensitivity, also showed substantial differences between MLH3 hotspot sites and the genome average (Supplemental Figure 4). However, there was no obvious difference in nucleosome occupancy levels between hotpot sites at the three prophase substages.

**Figure 6.**
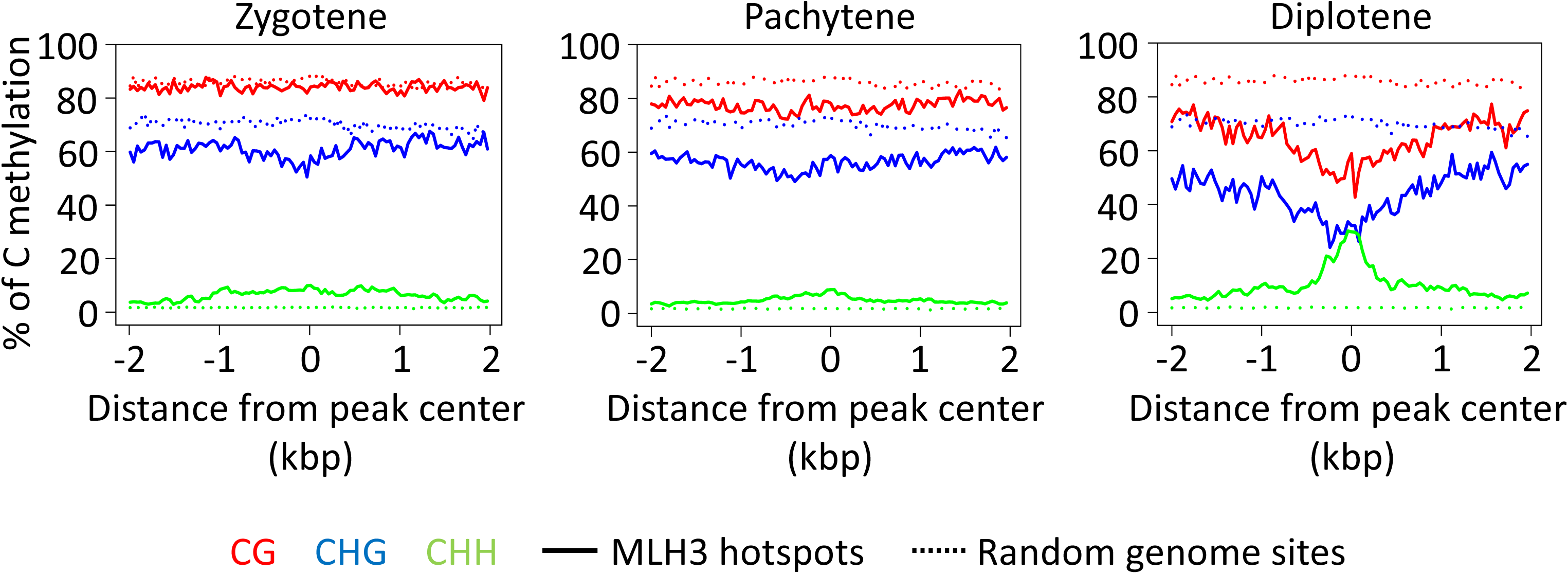
DNA methylation patterns at MLH3 hotspots at different substages of prophase I in the B73 maize inbred.

## DISCUSION

### High-resolution mapping of COs and CO intermediates identifies distinct transitions in CO formation

We developed a robust, high-resolution method to map COs and CO intermediates in meiosis, which allowed us to uncover novel features of CO locations and new aspects of the CO/non-CO decision. CO mapping using MLH3 ChIP is more precise, cheaper, and faster than conventional genetic mapping and is independent of the level and distribution of DNA sequence polymorphisms. MLH3 ChIP enables surveying millions of CO events, instead of mere thousands that can be examined with conventional genetic mapping. The relatively low CO number that can be analyzed cost-effectively with conventional methods is a major challenge of recombination studies in higher eukaryotes. It makes it difficult to identify CO hotspots and determine their activity. In maize, only a handful of CO hotspots have been known (Dooner and Martinez-Ferez, 1997; Yandeau-Nelson et al., 2006; Yao et al., 2002) and comprehensive CO hotspot studies have not been performed until now. In addition, MLH3 ChIP enables generating genetic maps in germplasm where conventional mapping is ineffective, such as inbreds and hybrids exhibiting limited inter-parental DNA sequence polymorphism. Using this approach, we found that a large proportion of COs in maize are formed in SNP desert regions, where conventional methods are unable to determine precise CO locations. MLH3 ChIP should also be useful for genetic mapping in species in which creating mapping populations is challenging and in inter-specific hybrids, where mature COs may lead to chromosome instabilities, preventing conventional mapping (Li et al., 2021; Morales and Dujon, 2012).

The ability of MLH3 ChIP to identify CO intermediates with high resolution allowed us to follow CO formation dynamics throughout prophase I. Interestingly, we found substantially fewer MLH3 hotspots in zygotene than in pachytene and diplotene. Chromosomal MLH3 foci were, however, more numerous in zygotene than in late prophase. These two observations imply that the distribution of MLH3 sites in zygotene is more uniform than in pachytene or diplotene. However, at most of these numerous initial sites there are insufficient MLH3 numbers at each site to make them hotspots. In contrast, MLH3 foci remaining by pachytene are at fewer but more active sites. The locations of diplotene MLH3 hotspots mostly overlap with the locations of pachytene hotspots. There is, however, a continuing decrease of MLH3 focus numbers from pachytene to diplotene. These data imply that there are two steps when MLH3 foci are “thinned down”. During the transition from zygotene to pachytene, thinning down involves non-CO-type resolution of dHJs at sites that are outside of CO hotspots. In contrast, the dHJs thinned down from pachytene to diplotene are already at CO hotspot sites.

It is intriguing that a large percentage of the strongest MLH3 hotspots are at the 180bp knob repeats. We believe, however, that this high frequency does not suggest that a disproportionate number of COs are located at the knobs but rather that knobs are consistently sites of COs in a high fraction of meioses. Interestingly, we did not find MLH3 hotspots associated with TR1 repeats, which are also present at various proportions at maize knobs (Ananiev et al., 1998; Ghaffari et al., 2013). The 180bp and TR1 repeats may convey distinct properties to knobs. Interestingly, the 180bp but not the TR1 repeats are involved in knobs acquiring neocentromere functions (Dawe et al., 2018).

### Recombination landscapes of hybrid and inbred maize are not significantly different

Using MLH3 ChIP, we were able to generate the first high-resolution recombination map of a maize inbred line. There have been numerous studies on the effect of inter-parental sequence polymorphism on recombination outcomes. Studies in plants have shown that higher CO rates associated with low as well as elevated levels of local DNA sequence polymorphism between parental chromosomes (Blackwell et al., 2020; Dooner, 2002; Emmanuel et al., 2006; Ziolkowski et al., 2015). However, the complex dynamics of meiotic chromosome interactions (Sheehan and Pawlowski, 2009) could potentially result in distant sequence polymorphisms having confounding effects on CO formation. Examining CO landscape in homozygous backgrounds enables complete elimination of inter-parental sequence polymorphism as a factor. Our results suggest that the effect of inter-parental DNA sequence polymorphism on CO landscape is negligible at the chromosome-wide scale and relatively minor at the fine scale. The residual effect of inter-parental polymorphisms on recombination outcomes may be related to the processes of dHJ formation and resolution, which are DNA sequence dependent (Cloud et al., 2012; Marsolier-Kergoat et al., 2018). It is also possible that the effect of inter-parental polymorphism is due to chromatin state differences between parental chromosomes instead of DNA sequence polymorphisms. DNA methylation patterns can largely persist in hybrids for several generations (Reinders et al., 2009). Thus, if parental lines display chromatin status differences, these differences are likely to be still present at subsequent meioses.

### Chromatin-dependent CO site designation

By performing MLH3 ChIP at different substages of prophase I, we were able to track CO intermediates and examine chromatin factors at their locations. We found that the change of MLH3 hotspot distribution from the DSB-like (He et al., 2017) in zygotene to the CO-like (Kianian et al., 2018) in late prophase was accompanied by distinct chromatin state characteristics at MLH3 hotspots at the different prophase substages. In particular, sites of MLH3 hotpots present in late prophase exhibited progressive reduction of CG and CHG methylation compared to hotspot sites in zygotene. These data fit into earlier observations that CO sites in maize exhibit severe reduction of CG and CHG methylation levels (Kianian et al., 2018) while at DSB hotspots this reduction is limited (He et al., 2017).

Our data suggest that reduced DNA methylation increases the likelihood of dHJ resolution as a CO. It is possible that such low-DNA-methylation chromatin environment is favored by the large protein complexes involved in CO formation, which require topologically-relaxed chromatin and unobstructed DNA access (Marsolier-Kergoat et al., 2018). However, since DNA demethylation during the course of prophase I is unlikely in plants (Zhang et al., 2018), there are only two scenarios that could result in reduced DNA methylation at MLH3 sites in late prophase: (i) resolution of dHJ at high-methylation sites into non-COs preceding CO formation and (ii) movement of MLH3-bound CO intermediates to sites of lower DNA methylation. Our data are consistent with both of these mechanisms. The reduction in the numbers of chromosomal MLH3 foci during the course of prophase I favors the former possibility while the reduction of the distance between MLH3 hotspots and CO sites during prophase progression favors the latter. Studies showing that MLH1/3 protein complexes can move along DNA and potentially operate as sliding clamps (Gallardo et al., 2015; London et al., 2021) gives additional credence to the latter hypothesis.

Altogether, our data imply a multi-step thinning down process of recombination intermediates during the progression of recombination. We show that early in the recombination pathway, the span of DSB resection is associated with a higher likelihood of a DSB being repaired as a COs. The final step of thinning down occurs during dHJ resolution and likely involves the MLH1/MLH3 protein complex. This step is responsible for the eventual shape of recombination landscape in maize, with most COs located towards chromosome arms and very few of them in pericentromeric regions. The extent of DNA methylation is critical for the outcome of this process. Future studies may address the specific nature of the interaction between MLH1/MLH3 and chromatin organization.

## MATERIALS AND METHODS

### Plant growth

B73 inbred and Mo17 x B73 hybrid maize plants were grown at the 12h/12h photoperiod, with 31°C, 70% air humidity and 500µmol/m^2^/sec^2^ light intensity during the day and 22°C and 70% humidity at night. At 42-50 days after planting, whole meiotic tassels were collected and analyzed to determine meiosis stage.

### Meiotic tassel staging

Several flowers from each tassel were fixed in 3:1 ethanol: acetic acid (v/v) solution for a minimum of one week. Anthers were dissected, used to prepare microscopic slides, and stained in 4ʹ,6-diamidino-2-phenylindole (DAPI) as previously described (Zelkowski and Pawlowski, 2020). Slides were evaluated using fluorescence microscopy.

### S1 nuclease mapping of DSB resection

To map DSB resection, we followed the protocol of Mimitou and Keeney (Mimitou and Keeney, 2018) with few modifications. Sixty zygotene anthers were collected and digested in 200µl of protoplasting enzyme mix (200mM sucrose, 10mM MES pH 5.2, 2mM CaCl_2_, 10mM KCl, 0.1% BSA, 2.5% cellulase, 1.0% macerozyme, 1.0 hemicellulase, 1% pectolyase) for 60min at 37°C. Next, the cells were embedded in 200µl of melted 2% low-melting agarose (Sigma-Aldrich, catalog No. 50111). The mixture was pippeted onto a sterile Petri dish to form a drop and moved to 4°C for 30min. Next, the agarose drop was cut into three equal pieces, each transferred to a separate 200µl Eppendorf tubes. The agarose plugs were treated with Solution 2 (0.45M EDTA pH 8, 0.01M Tris pH 7.5, 7.5% β-mecraptoethanol and 20µl of 100µg/ml RNAse C) and incubated for 1h at 37°C. Following incubation, the buffer was discarded. Solution 3 (0.25M EDTA pH 8, 0.01M TRIS pH 7.5, 1% SDS and 1mg/ml proteinase K) was applied, and samples were incubated overnight at 40°C. The following day, the agarose plugs were washed three times with 50mM EDTA pH 8.0 at room temperature, followed by three washes with 250µl of S1 nuclease buffer (Sigma-Aldrich, catalog No. M5761), for 45min each wash. Next, the plugs were incubated with 22µl of S1 nuclease (Sigma-Aldrich, catalog No. M5761) for 30min at 37°C. The S1 nuclease reactions were stopped by adding EDTA pH 8 to the final concentration of 10mM and placing on ice for 15min. The plugs were rinsed twice with 500µl of 1x TE buffer (10mM Tris-HCl, 0.1mM EDTA), followed by four washes with 250µl of T4 polymerase buffer (Sigma-Aldrich, catalog No. 101228-186). Next, 40µl of T4 polymerase (Sigma-Aldrich, catalog No. 101228-186) was added and the samples were incubated for 15min at 12°C. The reactions were stopped by adding EDTA pH 8 to the final concentration of 10mM and placing on ice for 15min, followed by a brief wash in 1x TE buffer (10mM Tris-HCl, 0.1mM EDTA). The remaining liquid was discarded and the tubes containing the agarose plugs were incubated at 75°C for 20min to inactivate the T4 polymerase. Then, the tubes were placed at room temperature for 10min, followed by 10min on ice, followed by four 10min washes in 250µl of T4 ligase buffer. Next, 1000 units of T4 ligase and 50mM of biotinylated adapters were added. The reactions were performed overnight at 16°C. Then, the plugs were washed with 500µl buffer containing 5mM Tris-HCl pH 7.5, 0.5mM EDTA and 1M NaCl, followed by two washes in 10mM Tris–HCl pH 7.5. Biotinylated DNA fragments were purified following the NucleoSpin Gel clean-up kit (Sigma-Aldrich, catalog No. 740609.50) protocol. Next, the DNA fragments were bound to M-280 Streptavidin Dynabeads™ (Sigma-Aldrich, catalog No. 11205D) following the manufacturer’s protocol.

### Chromatin immunoprecipitation using MLH3 antibodies

Chromatin immunoprecipitation experiment was performed as previously described (He et al., 2017). Tassels at desired meiosis stages were fixed in a 1% freshly made formaldehyde solution containing 10mM Tris HCl pH=8.0, 0.4M sucrose, 10mM MgCL_2_, and 5mM β-mercaptoethanol under vacuum at room temperature for 20min. The fixation reaction was stopped by adding glycine to the final concentration 0.125M. Fixed flowers were briefly dried on filter paper, flash frozen in liquid nitrogen, and stored at -80°C. Flowers collected from 4 to 8 tassels were used for each ChIP experiment. Tissue was ground to fine powder in liquid nitrogen, homogenized in extraction buffer A (10mM Tris-HCl, 0.4M sucrose, 10mM MgCl_2_, 2mM PMSF, 5mM β-mercaptoethanol, 1µg/ml protease inhibitor (Sigma-Aldrich, catalog No. 200-664-3), incubated with gentle shaking for 20min, filtered through miracloth, and centrifuged at 11 000rpm at 4°C for 20min. The resulting pellets were resuspended in extraction buffer B (10mM Tris-HC, 0.25M sucrose, 10mM MgCl2, 1% Triton x-100, 2mM PMSF, 5mM β-mercaptoethanol, 1µg/ml protease inhibitor (Sigma-Aldrich, catalog No. 200-664-3). Next, the solutions were centrifuged at 14 000rpm for 10min at 4°C. Pellets were resuspended in extraction buffer C (10mM Tris-HCl, pH=8, 1.7M sucrose, 2mM MgCl_2_, 0.15% Triton X-100, 2mM PMSF, 5mM β-mercaptoethanol, 1µg/ml protease inhibitor (Sigma-Aldrich, catalog No. 200-664-3) and centrifuged at 14 000rpm for 1h at 4°C. Pellets were resuspended in 620µl of nuclei lysis buffer (50mM Tris-HCl, 10mM EDTA, 1% SDS, 2mM PMSF, 2µg/ml protease inhibitor (Sigma-Aldrich, catalog No. 200-664-3). The suspensions were sonicated for 20min with the Bioruptor® Sonication System (Diagenode, catalog No. UCD-200) using a 30sec high and 30sec break setting. Chromatin samples were split into six ChIP samples: 2 replicates for MLH3-N and MLH3-C antibodies each and one replicate each for MLH3-N and MLH3-N pre-immune sera. Samples were diluted by adding 300µl of ChIP dilution buffer (16.7mM Tris-HC, 1.2mM EDTA, 167mM NaCl, 1.1% Triton X-100, 1mM PMSF, 5mM β-mercaptoethanol, 1µg/ml protease inhibitor (Sigma-Aldrich, catalog No. 200-664-3).

Protein A-coated Dynabeads™ (Invitrogen, catalog No. 10001D) were incubated with BSA to the final concentration of 2.5 mg/ml with gentle shaking for 2h. Chromatin samples were precleared by mixing with the beads for 2h at 4°C. After pre-clearing, beads were collected on a magnetic separation stand and supernatants were transferred into new tubes. 10µg of MLH3-C or MLH3-N antibodies were added to each -pre-cleared sample. For negative controls, 40µg of pre-immune serum was added to each tube. The samples were placed overnight at 4°C with gentle shaking. Next day, 45µl of BSA coated protein A Dynabeads™ Protein A were added to each sample and the samples were placed for 2h at 4°C with gentle shaking. Subsequently, the tubes were placed on a magnetic separation stand, supernatants were discarded and beads were washed with Low Salt Wash Buffer (20mM Tris HCl pH=8, 2mM EDTA, 150mM NaCl, 0.05% SDS, 1% Triton X-100) for 10min at room temperature with gentle shaking. Tubes were placed on a magnetic separation stand, supernatant were discarded, and beads were washed with High Salt Wash Buffer (20mM Tris HCl pH=8, 2mM EDTA, 500mM NaCl, 0.05% SDS, 1% Triton X-100) for 10min at room temperature with gentle shaking. Tubes were placed on a magnetic separation stand, supernatants were discarded, and beads were washed with LiCl Wash Buffer (10mM Tris HCl pH=8, 1mM EDTA, 250mM LiCl, 1% NP-40, 0.5% sodium deoxycholate) for 10min at room temperature with gentle shaking. Tubes were placed on a magnetic separation stand, supernatants were discarded, and beads were washed twice with TE buffer (10mM Tris HCl pH=8, 1mM EDTA) for 10min at room temperature with gentle shaking. Tubes were placed on a magnetic separation stand, supernatants were discarded, and chromatin fragments were eluted using 100µl of Elution Buffer (50mM Tris-HCl pH=8.0, 10mM EDTA, 1% SDS, 200mM NaCl) at 65°C for 30min. Elution was performed twice. Eluates were combined and decrosslinked in Elution buffer at 65°C for 8h. Next, the sample were treated with RNase at the final concentration of 100µg/ml for 2h at 37°C. Finally, the samples were digested using proteinase K at the final concentration of 200µg/ml for 2h at 42°C.

### DNA purification

To recover DNA from S1 nuclease mapping and ChIP experiments, glycogen to the final concentration of 0.5µg/ml and 1/10 of sample volume of 3M sodium acetate were added to each DNA sample, followed by vortexing. Next, an equal volume of isopropanol was added, mixed gently, and the samples were incubated for 1h at -20°C. The samples were centrifuged in 4°C at 12,000 x g for 15-30min, supernatants were discarded, and pellets were washed with cold 70% ethanol. The samples were then centrifuged in 4°C at 12,000 x g for 15-30 min, supernatants were discarded, and pellets were air dried for 20-30 min. The pellet was dissolved in 50-100µl of TE buffer.

### Illumina library construction

DNA fragments recovered from S1 nuclease mapping and ChIP experiments were used to construct Illumina sequencing libraries using the NEBNext® Ultra™ II DNA Library Prep Kit for Illumina® (NEB, catalog No. E7645L) NEBNext® Multiplex Oligos for Illumina® Index Primer sets 1, 2, and 3 (NEB, catalog No. E7500S) following manufacturer’s instructions. The libraries were amplified using 15 PCR cycles, purified as described above, and sequenced using Illumina NextSeq 500 with 75bp sequencing length.

### Immunolabeling of maize meiocytes

Immunolabeling was performed as previously described (Zelkowski and Pawlowski, 2020). Maize tassels at appropriate meiosis stages were placed in Carnoy’s fixative (3:1 v/v ethanol : acetic acid) and incubated overnight at room temperature. Then, the fixative was replaced with a fresh aliquot and fixation was continued for at least four more days. Anthers were dissected into small Petri dishes, washed three times with 3ml of Carnoy’s fixative, and placed in ice-cold 10mM citric buffer pH 4.6. The anthers were then washed three times with 10mM citric buffer pH 4.6 and digested in 1ml of enzyme mix (0.1g Onozuka R10 Cellulase (Grainger), 0.1g cytohelicase (Sigma), 0.1g pectolyase from *Aspergillus japonicus* (Sigma) in 29 ml of 10mM citric buffer, pH 4.6) for 120min at 37°C. Digestion was stopped by adding 2 ml of ice-cold 10mM citric buffer, pH 4.6.

To make microscopic slides, single anthers were placed on glass slides and macerated with metal rod to release meiocytes. Then, 10µl of 60% ice-cold acetic acid was added. The slides were placed on a warm plate set at 43°C for 50sec, followed by addition of another 10µl of ice-cold 60% acetic acid. Next, 40µl of ice-cold Carnoy’s fixative was used to form a ring around the meiocyte drop and the slide was rinsed with 200µl of Carnoy’s fixative. The slides were then air dried for 10min, followed by a quick wash in 10mM citric acid buffer pH 6.0. The slides were then microwaved for 35sec and placed in cold TBST (137mM NaCl, 12mM Phosphate, 2.7mM KCl, 1% Tween-20 pH 7.4).

For immunolabelling, slides were washed three times with TBST and air dried for 30sec. Then, 50µl of blocking solution (1% bovine serum albumin in TBST) was added and incubated for 30 min at room temperature. Next, slides were washed three times in TBST and air dried for 1min. Then, the slides were incubated with the primary antibody solution at 4°C overnight. Next, the slides were washed three times with TBST and incubated with secondary antibody at room temperature for 90min. Finally, the slides were washed three times with TBST, air dried for 90sec, and mounted in DAPI-containing Vectashield (Vector laboratories). Slides were analyzed under a fluorescence microscope.

### Mapping Illumina sequence reads

Illumina sequence reads were mapped onto the maize reference genome (Jiao et al., 2017) using bowtie2 (Langmead et al., 2009) version 2.4.3 with the argument “--local” on. Reads with mapping quality (MQ) of 5 or less were discarded. Samtools’s “rmdup” command was used to remove duplicate reads with default settings, followed by “sort”, “index”, and “output BAM format”.

### Calling of S1 and MLH3 hotspots

The two independent replicates were merged using the merge command of Samtools (Li et al., 2009). Enrichment regions were defined with MACS2 (version 2.2.7.1) (Zhang et al., 2008), and the subcommand callpeak was used with the following parameters -g 2.0e^9^ --nomodel --extsize 300 --keep-dup=auto -m 2 5 and the FDR cutoff of 0.05. The genome input was used as control.

### DNA methylation analysis

Previously published bisulfate sequencing data (He et al., 2017) were used for the DNA methylation analyses. Illumina reads were trimmed to remove adapters, filtered to remove poor-quality reads, and mapped to the reference genome using Bismark (Krueger and Andrews, 2011) with default setting. Percentage of methylated cytosines in the CG, CHG or CHH contexts were calculated at MLH3 hotspot sites as 100* (methylated reads)/ (methylated reads + unmethylated reads)

### Nucleosome occupancy analysis

Previously published data (He et al., 2017) were used for nucleosome occupancy mapping. Illumina reads were mapped to the maize reference genome (Jiao, 2017) using bowtie2 (Langmead, 2009) version 2.4.3 with the argument “--local” on. Reads with mapping quality (MQ) of 5 or less were discarded. Deeptools bamCoverage (https://deeptools.readthedocs.io/en/develop/) was used to generate bigwig files with bin size of 200bp normalize to read count per bin level. ComputeMatrix and plotProfile were used for plotting nucleosome occupancy at MLH3 hotspot sites.

### MLH3 hotspot analyses

To identify the proportion of MLH3 hotspots overlapping genetically mapped maize COs, we calculated the proportion of MLH3 hotspots, defined as windows centered on hotspot peaks, that overlapped midpoints of COs genetically detected in the NAM and B73 x Mo17 populations (Kianian et al., 2018; McMullen et al., 2009). DNA sequence motifs associated with MLH3 hotspots were determined with meme (Bailey et al., 2015) using the range of 30 nucleotides and default settings. To identify MLH3 hotspots located in the 180bp knob repeats, sequences from GenBank accessions AF030934.1, M32521.1, M32525.1, DQ352544.1 and DQ352544.1 containing the 180bp knob repeat sequence were used to construct a PWM matrix with meme (Bailey et al., 2015). The matrix was then used to identify knob regions in the maize genome. Only hits with E-values lower than 1e^-50^ were used.

### Statistical analyses

All statistical analyses were performed with the R v.4.1.2 software.

### Accession numbers

The S1 mapping and MLH3 ChIP sequence data reported in this paper have been deposited in the National Center for Biotechnology Information (NCBI) Gene Expression Omnibus (GEO) database, accession number GSE210615.

## ACKNOWDEGEMENTS

We would like to thank Kasia Zelkowska and Joshua Derrick for expert technical assistance. This work was supported by an NSF grant IOS-1546792 to WPP. W.P.P. is a Cornell Institute for Food Systems Faculty Fellow.

