## Supplemental Tables and Figures for "Crossing-over decision landscape in maize"

Supplemental Table 1. Numbers of unique reads aligning to the maize genome sequence scaffold in S1-based mapping of DSB resection.

| **Sample** | **Number of unique mapping reads** |
| --- | --- |
| S1-treated B73 - replicate 1 | 6804896 |
| S1-treated B73 - replicate 2 | 11949874 |
| B73 input DNA | 14639531 |
| Untreated B73 control | 1788185 |
| S1-treated B73 leaf (somatic cells) | 6766 |
| S1-treated *ameiotic1* | 5168 |

Supplemental Table 2. Spearman correlation between samples in S1-based mapping of DSB resection.

| | S1-treated B73 - replicate 1 | S1-treated B73 - replicate 2 | Untreated B73 control | B73 input DNA |
|---|---|---|---|---|
| **S1-treated B73 - replicate 1** | 1 | 0.93 | 0.75 | 0.52 |
| **S1-treated B73 - replicate 2** | | 1 | 0.77 | 0.54 |
| **Untreated B73 control** | | | 1 | 0.47 |
| **B73 input DNA** | | | | 1 |

Supplemental Table 3. Numbers of unique reads mapping to the maize genome in MLH3 ChIP experiments.

| **Genotype** | **Stage** | **Antibody** | **Numbers of unique sequence reads (mln)** |
|---|---|---|---|
| B73 x Mo17 | zygotene | MLH3-C | 29 |
| B73 x Mo17 | zygotene | MLH3-N | 29 |
| B73 x Mo17 | pachytene | MLH3-C | 64 |
| B73 x Mo17 | pachytene | MLH3-N | 64 |
| B73 x Mo17 | diplotene | MLH3-C | 27 |
| B73 x Mo17 | diplotene | MLH3-N | 47 |
| B73 | zygotene | MLH3-C | 73 |
| B73 | zygotene | MLH3-N | 22 |
| B73 | pachytene | MLH3-C | 10 |
| B73 | pachytene | MLH3-N | 18 |
| B73 | diplotene | MLH3-C | 18 |
| B73 | diplotene | MLH3-N | 40 |

Supplemental Table 4. Numbers of MLH3 hotspots identified in the B73 x Mo17 hybrid colocalizing in experiments using antibodies targeting the C and N termini of MLH3. Window size = 2kb; *q* = 0.05.

| **Stage** | **Peak No. for MLH3-C** | **Peak No. for MLH3-N** | **No. of overlapping peaks** | **% of overlapping peaks** |
|---|---|---|---|---|
| Zygotene | 61 | 690 | 48 | 78.7 |
| Pachytene | 1115 | 1066 | 673 | 63.1 |
| Diplotene | 6945 | 63426 | 5503 | 79.2 |

Supplemental Table 5. Correlation between MLH3 patterns generated using antibodies targeting the C and N termini of MLH3 at different prophase I substages in the B73 x Mo17 hybrid. Table shows numbers of MLH3 peaks colocalizing in non-overlapping 2 kb windows distributed genome wide. $q$ = 0.05.

| **Pattern 1** | | | ***vs.*** | **Pattern 2** | | | **No. of overlapping peaks** | **% of overlapping peaks** |
|---|---|---|---|---|---|---|---|---|
| **Stage** | **Antibody** | **Peak No.** | | **Stage** | **Antibody** | **Peak No.** | | |
| zygotene | MLH3-C | 61 | | pachytene | MLH3-C | 1115 | 9 | 14.8 |
| zygotene | MLH3-C | 61 | | diplotene | MLH3-C | 6945 | 17 | 27.9 |
| pachytene | MLH3-C | 1115 | | diplotene | MLH3-C | 6945 | 974 | 87.4 |
| zygotene | MLH3-N | 690 | | pachytene | MLH3-N | 1066 | 14 | 2.0 |
| zygotene | MLH3-N | 690 | | diplotene | MLH3-N | 63426 | 92 | 13.3 |
| pachytene | MLH3-N | 1066 | | diplotene | MLH3-N | 63426 | 963 | 90.3 |

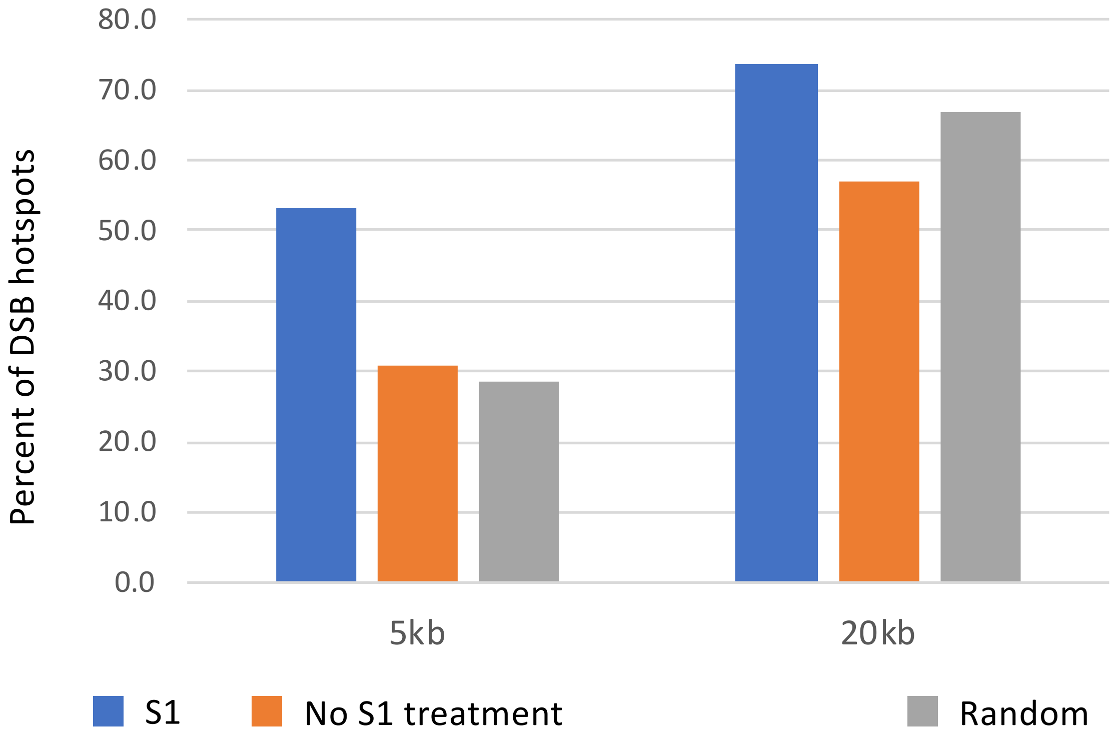


**Supplemental Figure 1.** Percent of DSB hotpots identified using RAD51 ChIP located within 5kb and 20kb from S1 resection sites 500bp to 2000bp in length.

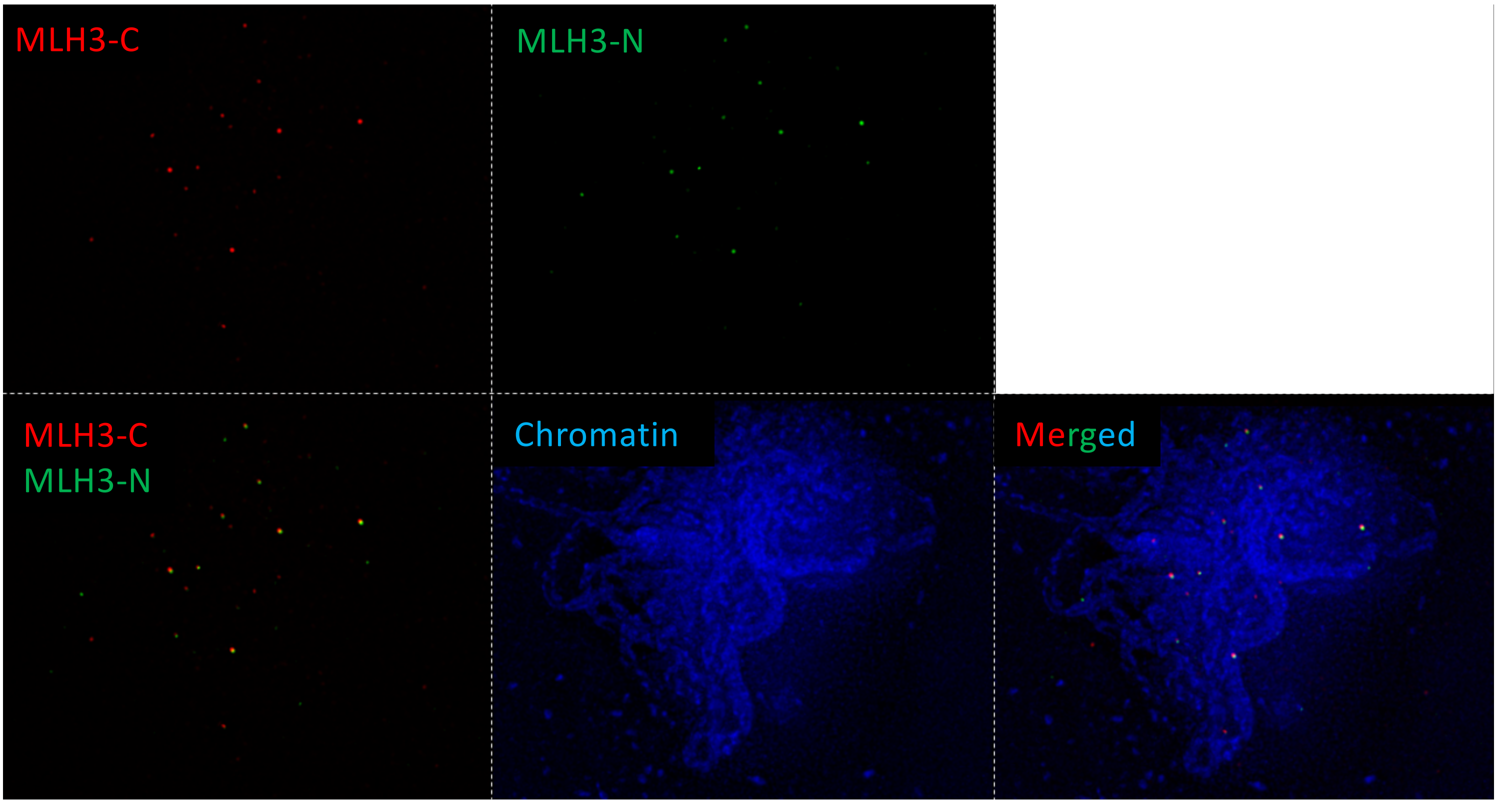


**Supplementary Figure 2.** Co-immunolabeling of MLH3-C and MLH3-N antibodies pachytene chromosomes in the B73 x Mo17 hybrid.

A

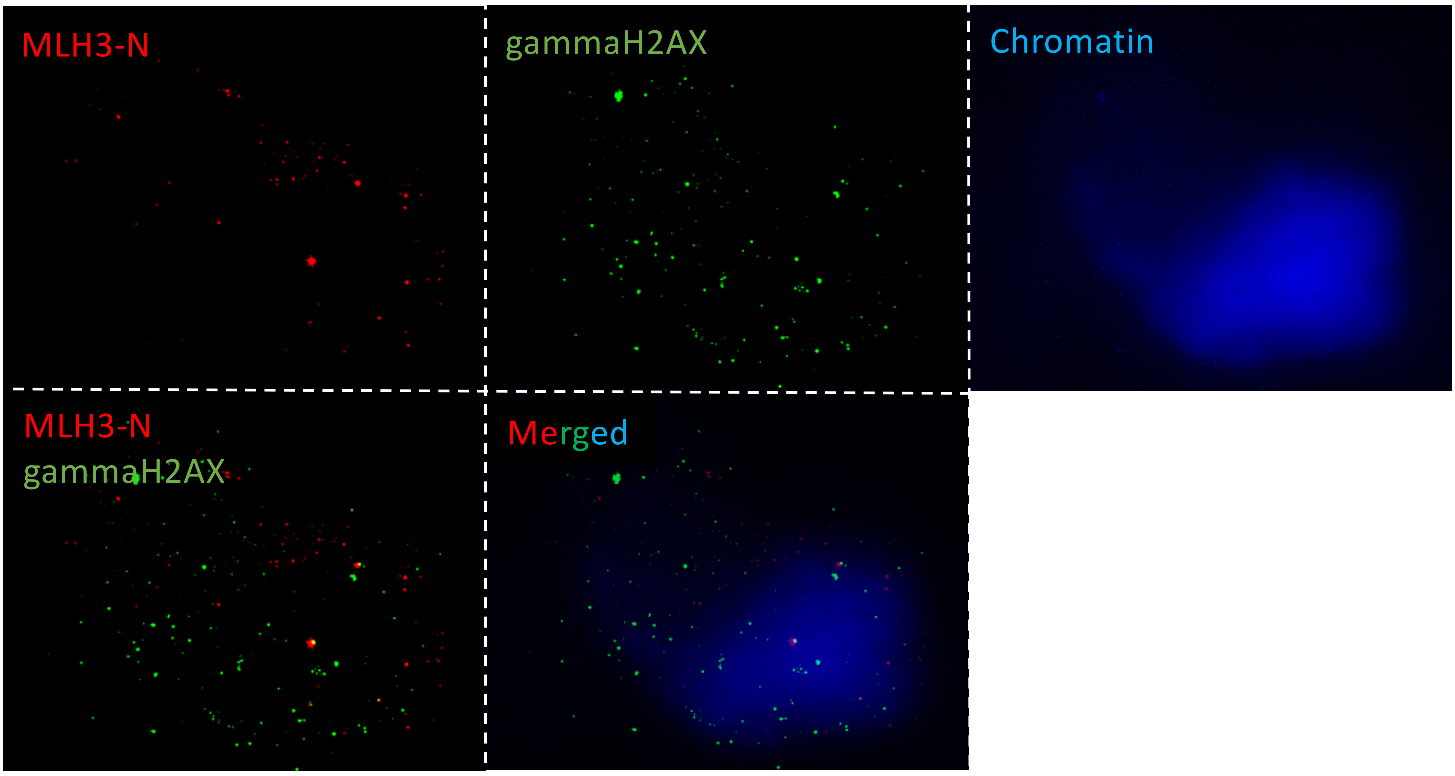


B

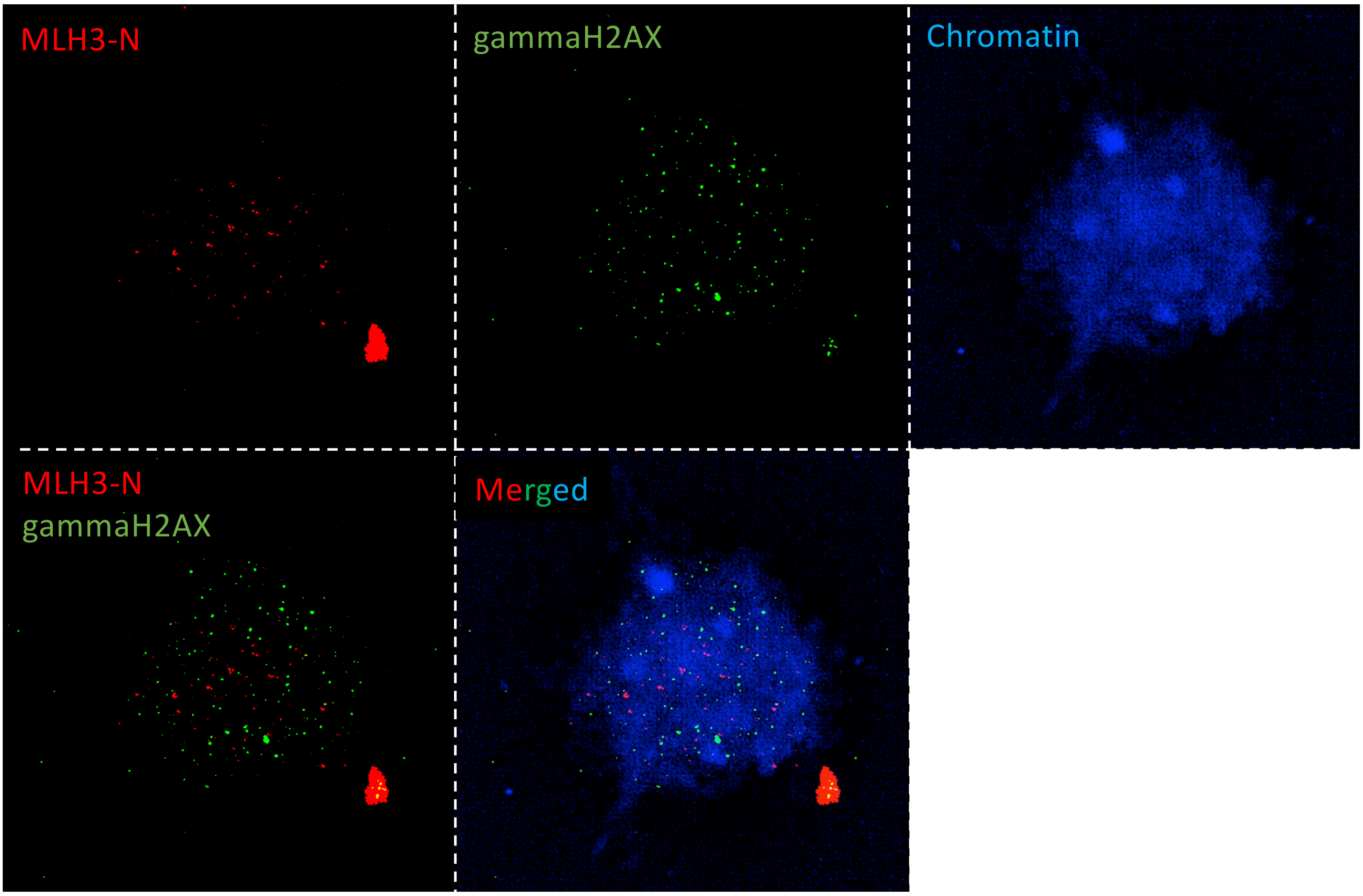


**Supplementary Figure 3.** Co-localization of MLH3-N foci with early recombination markers gammaH2AX (A) and RAD51 (B).

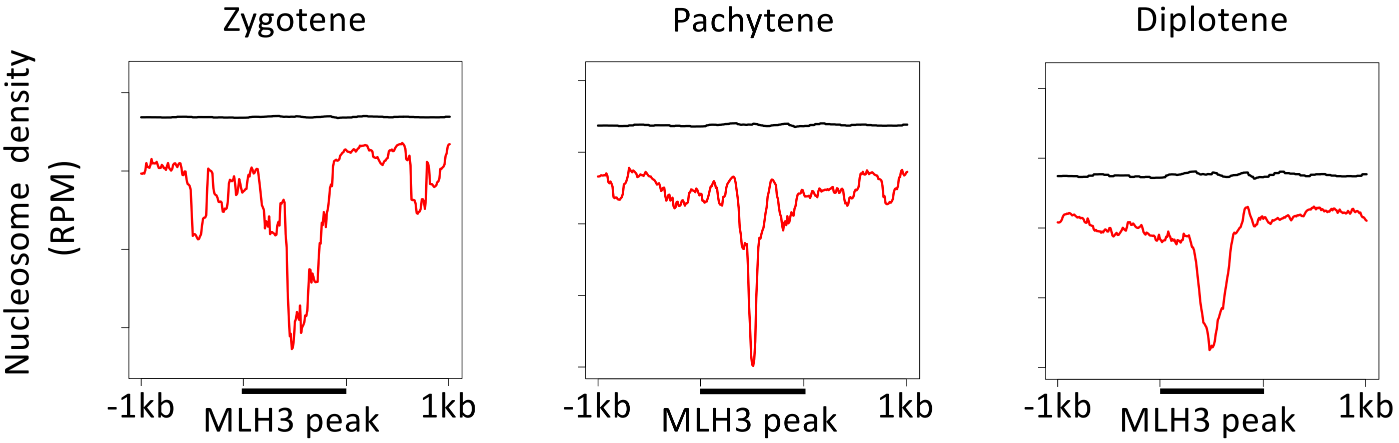


**Supplementary Figure 4.** Nucleosome occupancy measured by monococcal nuclease digestion at the sites of MLH3 hotspots detected at different substages of meiosis I in the maize inbred B73.
